# Male pathology regardless of behaviour drives transmission in an avian host-pathogen system

**DOI:** 10.1101/2023.09.28.559954

**Authors:** Erin L. Sauer, Chloe Connelly, Weston Perrine, Ashely C. Love, Sarah E. DuRant

## Abstract

Host sex is an important source of heterogeneity in the severity of epidemics. Pinpointing the mechanisms causing this heterogeneity can be difficult because differences in behaviour among sexes (e.g. greater territorial aggression in males) can bias exposure risk, obfuscating the role of immune function, which can lead to differences in pathology, in driving differential susceptibility between sexes. Thus, sex-biased transmission driven by differences in immune function independent of behaviour is poorly understood, especially in non-mammalian systems. Here we examine the previously unexplored potential for male-biased pathology to affect transmission using an avian host-pathogen system. We employ a sex-dependent multistate transmission model parameterized with isolated, individual-based experimental exposures of domestic canaries and experimental transmission data of house finches. The experiment revealed that male birds have shorter incubation periods, longer recovery periods, higher pathogen burdens, and greater disease pathology, than females. Our model revealed that male-biased pathology led to epidemic size rapidly increasing with the proportion of male birds, with a nearly 16-fold increase in total epidemic size from an all-female to an all-male simulation. Our results demonstrate that female-biased resistance, independent of male behaviour, can drive sex-dependent transmission in wildlife, indicating that sex-based differences in immune function, not just differences in exposure risk, can shape epidemic dynamics.

## Introduction

Heterogeneity in exposure risk (*the likelihood of being exposed to a pathogen*), resistance (*ability to prevent or limit pathogen growth*), and pathology (*the intensity of symptomatic disease*) can have major impacts on transmission dynamics (Paull et al., 2012). Transmission estimates that fail to account for this heterogeneity can result in poor estimates of disease dynamics and ineffective disease control measures (Lloyd-Smith et al., 2005). One such driver of heterogeneity is host sex. Sex-based differences in behavior (e.g., frequency and types of social interactions) and physiology contribute to differences in exposure risk, disease susceptibility and pathology, and transmission (Grear et al., 2009; Hawkins et al., 2006; McCurdy et al., 1998; Mougeot et al., 2005; Nielsen et al., 2020; Perkins et al., 2008; Zuk & McKean, 1996).

Some sex-biased behavioural and physiological traits contributing to host heterogeneity are intertwined, potentially resulting in additive or multiplicative effects on disease transmission that can be hard to disentangle. For example, in vertebrate systems, males are generally thought to be more likely to become infected and harbor larger infection burdens than females (Zuk & McKean, 1996). Male-biased territorial behaviours can result in males having larger range sizes, contact networks, and aggressive encounters, thus increasing their exposure and transmission risks (Grear et al., 2009; Hawkins et al., 2006; Miller & Conner, 2005). Simultaneously, the propensity for territorial behaviour and aggression in males is often associated with increased stress and testosterone, both of which can be immunosuppressive (Duckworth et al., 2001; Ferrari et al., 2004; Grear et al., 2009; Hawley, 2006; Love et al., 2017; Mougeot et al., 2005; Zuk & McKean, 1996). Thus, the role of immune function, independent of host behaviour, in sex-biased infection remains unclear as it is often difficult to distinguish between differential exposure and differential susceptibility (Zuk & McKean, 1996).

Additionally, much of the evidence supporting male-biased transmission is focused in mammalian-macroparasite systems while less is known about microparasite infections and infections of non-mammalian hosts (Valdebenito et al., 2021). Recent meta-analyses of sex-biases in immune function across multiple taxa and specifically in avian hosts found weak to no evidence of female-biased immunocompetence and some evidence of male-biased immunocompetence in avian hosts (Kelly et al., 2018; Valdebenito et al., 2021). For example, while male-biased infection prevalence and burden has been demonstrated in avian hosts, female-biased infections are seemingly as common (Duckworth et al., 2001; McCurdy et al., 1998; Mougeot et al., 2005; Nielsen et al., 2020; Valdebenito et al., 2021). Further, few studies attempt to translate these sex-biases in infection to sex-biased transmission dynamics (Kulkarni & Heeb, 2007; Lachish et al., 2011). Thus, despite the generality with which male-biased transmission is often thought to occur, there is little evidence in avian pathogen systems to suggest that males disproportionally contribute to transmission.

Here we examine the potential for male-biased pathology, the intensity of symptomatic disease, to drive transmission dynamics of a widespread avian bacterial pathogen, *Mycoplasma gallisepticum* (hereafter MG). MG is spread through fomites and direct contact with infected birds, primarily house finches (*Haemorhous mexicanus*), with bird feeders being important drivers of transmission in this system (Adelman et al., 2015; Dhondt et al., 2007; Hochachka et al., 2021; Hochachka & Dhondt, 2000; Moyers et al., 2018). Because of the importance of bird feeders in this system, sex-specific feeding behaviour may contribute to sex-biased exposure risk (the likelihood of being exposed to MG). For example, Bouwman & Hawley (2010) demonstrated that healthy house finch males tend to preferentially feed near diseased-males to avoid costly aggressive encounters with other healthy-males while healthy-females show no feeding preference (Hawley, 2006; Hawley et al., 2006). Thus, while there is some indication that males have the potential to disproportionately contribute to transmission in this system, direct assessment of sex-biased transmission of MG remains underexplored (Adelman et al., 2015; Moyers et al., 2018).

The potential for male-biased pathology to affect recovery time and pathogen burdens, both important transmission factors, paired with male-biased behavioural feeding preferences that could increase exposure risk during epidemics, may result in drastically male-biased transmission in MG epidemics. Here we examine the potential for male-biased sex-dependent transmission in MG using a multistate transmission model parameterized with exposure-controlled pathological data from an original experiment using domestic canaries and transmission rates from published data on house finch flocks of varying sex-ratios (Adelman et al., 2015; Moyers et al., 2018). Our objectives were to determine if male-biased transmission occurs in this system and, if so, can the bias be attributed to male-biased disease pathology or behaviour. We predicted that transmission will primarily be driven by male birds and that male pathology will be an important driver of transmission.

## Materials and Methods

### Study species and animal husbandry

MG can infect numerous songbirds species, though it is best studied in house finches (Farmer et al., 2005). We used domestic canaries (*Serinus canaria domestica*) in this experiment as a model for the MG-house finch system because canaries and house finches do not differ in their pathogen loads, production of pathogen-specific serum antibodies, pathology, or recovery time after exposure to identical doses of MG (Hawley et al., 2011). Further, canaries are well suited for laboratory studies and continue to exhibit typical life history events in captivity. Additionally, while there is limited data available on sex-biased differences in MG pathology, we were able to find three studies that found sex-biased pathology in house finches that is similar to our findings in canaries (Adelman et al., 2015; Hawley et al., 2007; Moyers et al., 2018). However, this pattern is inconsistent as there were many that found no difference between sexes (Sydenstricker et al., 2006; Thomason et al., 2017; Vinkler et al., 2018; Weitzman et al., 2021) and a couple that found higher female pathology in house finches (Altizer et al., 2004; Leon & Hawley, 2017).

Canaries were housed individually in cages with ad libitum access to water and food on a 12L:12D light cycle. Exposed (*N* = 13; 6 female, 7 male) and sham-exposed individuals (*N* = 12; 6 female, 6 male) were housed on separate racks separated by an opaque room divider to prevent exposure to the pathogen or disease-related social cues (Love et al., 2021). Sexes were randomly distributed within racks. On day 0, birds were inoculated in the palpebral conjunctiva of both eyes with either 0.025 mL of MG suspended in Frey’s media (5.00×10^7^ CCU/mL; VA1994; E. Tulman, University of Connecticut) or with a sham of Frey’s media alone. Canary experimental methods were approved by the University of Arkansas International Animal Care and Use Committee.

### Host responses

Disease pathology in canaries was assessed using conjunctival inflammation or “eye score” (0-3 scale per eye, summed to get “total eye score”; modified from Sydenstricker et al., 2006), body mass (g), and furcular fat score (0-3 scale in 0.5 intervals). Eye scores were recorded on days 0, 1, 3, 5, 7, then twice a week until day 35. Body mass and fat scores were recorded prior to exposure and on days 7, 14, 21, and 35. Blood samples and eye swabs were collected prior to exposure and on days 7, 14, and 21. During swabbing, a sterile swab was twirled along the conjunctiva of each eye for 5 sec and stored at -20°C. MG DNA was extracted using Qiagen DNeasy Blood and Tissue protocol (Qiagen, Inc., Valencia, CA) and quantified to determine pathogen load using qPCR methods based on Grodio *et al*. (2008) with gBlock® plasmid-based standards (Integrated DNA Technologies, Skokie, IL) described in Adelman *et al*. (2013). All birds were confirmed to be uninfected before the start of the experiment and sham-exposed birds never developed symptoms or tested positive for MG.

Blood samples were used to measure MG-specific antibody response, hematocrit, and differential leukocyte counts. Plasma was separated from whole blood after extraction and stored at -20°C. MG-specific IgY titers were quantified using an enzyme-linked immunosorbent assay (ELISA) kit (FlockChek *M. gallisepticum* ELISA kit, IDEXX, Westbrook, ME) with minor modifications following Love *et al*. (2021). Relative abundance of leukocytes were quantified blind of treatment from the feathered edge of blood smears (JorVet Dip Quick Stain Kin, Jorgensen Labs, Loveland, CO) by classifying the first 100 white blood cells observed (basophil, eosinophil, heterophil, lymphocyte, & monocyte). Relative abundance was quantified prior to and one week post exposure and the difference in pre-to post-abundance were calculated and compared (see statistical analysis). The canary experimental exposures and sampling timelines described above were designed for an unrelated study which is described in the Supplemental Methods.

### Statistical analysis

All statistics were conducted with R version 4.1.0 in R Studio (RStudio Team, 2021). To test for the effect of sex on total eye score, we conducted a Poisson distributed generalized additive mixed model that included a smoothing spline to model non-linear effects over time (*mgcv* package) (Preston & Sauer, 2020; Wood, 2011). To test for the effect of sex and MG exposure on body fat, mass (g), log_10_-transformed pathogen load, and hematocrit (%) over time we conducted linear mixed-effects models followed by ANOVAs that included main and interactive effects of sex, MG exposure, and days since exposure (*lme4 & car* packages) (Bates et al., 2014). To test for the effect of sex and MG exposure on change in leukocyte relative abundance from pre-exposure to one week post exposure, we conducted a MANOVA that included main and interactive effects of sex and MG exposure (*stats* package). We conducted an additional MANOVA to test for differences in leukocyte relative abundance between males and females prior to exposure. All repeated measures models included individual as a random intercept. Eye score, log_10_-transformed pathogen load, and antibody models were restricted to only MG-exposed individuals as no control birds ever developed symptoms or tested positive for MG or MG-specific antibodies. All other models included main and interactive effects of MG exposure, sex, and days since exposure. Model prediction plots were made using the predict function and the original conditions of the datasets.

### SEIR model

All simulations were conducted with R version 4.1.0 in R Studio using the *deSolve* package (RStudio Team, 2021; Soetaert et al., 2010). We created a mathematical model to determine the sex-dependent transmission dynamics of MG. The model subdivides the total population, *N(t)*, through time *t* by sex *i* and into susceptible, *S*_*i*_*(t)*, exposed, *E*_*i*_*(t)*, infected, *I*_*i*_*(t)*, recovered, *R*_*i*_*(t)*, and mortality, *M*_*i*_*(t)*, states (Figure 1) (Li & Muldowney, 1995). We ran 5 simulations of our model, varying the initial proportion of susceptible individuals that were male from 0-100% in 25% increments. Starting conditions were *S*=99, *E*=0, *I*=1, *R*=0, *M*=0. In all models with male individuals, the single infected bird in the starting conditions was male. We ran additional simulations where the index bird was female, and the results were nearly identical. Our model assumes density-dependent transmission with homogenous mixing of adult individuals, no birth or immigration, and that individuals who recover are no longer susceptible (see Supplemental Methods and Results for simulations based on frequency-dependent transmission). We intentionally excluded seasonal breeding dynamics from the model as parameters were measured from non-breeding birds and birth rates lack meaning in simulations with drastic sex ratio manipulations. Further, epidemics are largest during non-breeding making non-breeding individuals the most appropriate to model (Hosseini et al., 2004). Transmission dynamics are given by the system of differential equations:

**Figure 1.**
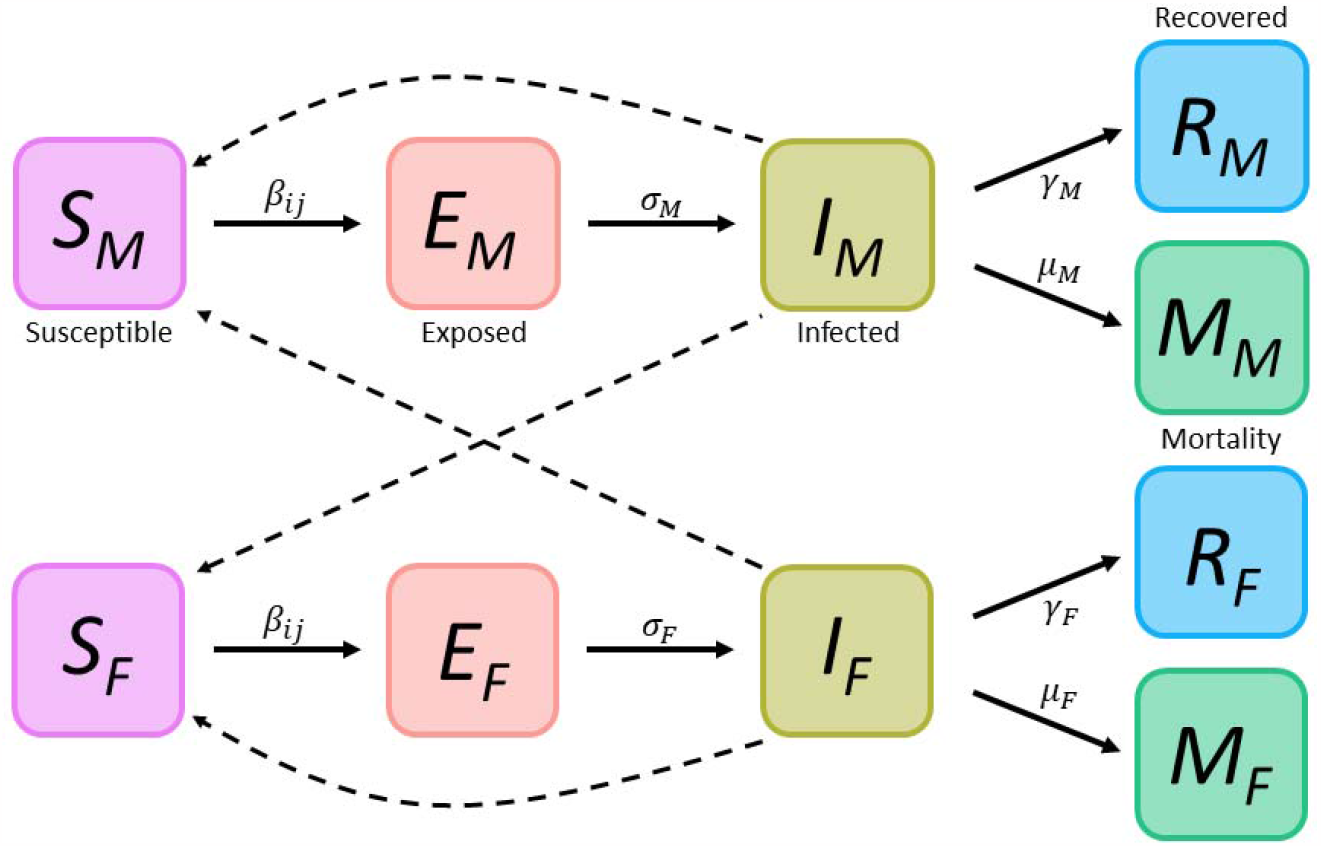
Scheme for the Susceptible, Exposed, Infected, Removed (recovered or mortality) transmission model for adult male (M) and female (F) *Mycoplasma gallisepticum* house finches (*Haemorhous mexicanus*).

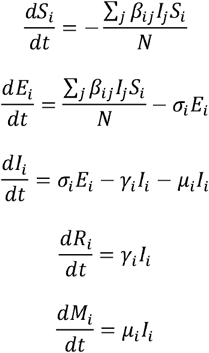

### Parameter estimates

Transmission rate from infectious sex *i* to susceptible host type *j* (*β*_*ij*_) was estimated from two published house finch – MG feeder transmission datasets (Adelman et al., 2015; Moyers et al., 2018) and is given by: 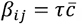, where *τ* is the prevalence of new infections among susceptible individuals and 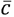 is the contact rate or the number of individuals in the flock divided by the duration of time since the last sampling period (Anderson & May, 1986). We use density-dependent transmission rates, as opposed to frequency-dependent, because density-dependence best describes transmission at feeders (Hosseini et al., 2004). Final *β*_*ij*_ values are the average of multiple same-type flock-level transmission rates. An individual was considered infected when MG load was > 100 gene copies (see Supplemental Methods & Table S1 for more details). Single-sex transmission rates were estimated from experimental same-sex house finch flocks published in Adelman et al. (2015) and included 6 male flocks and 4 female flocks sampled 3 times at 4 day intervals. Transmission rates from male and female birds to 50:50 mixed-sex flocks were estimated from experimental mixed-sex house finch flocks published in Moyers *et al*. (2018) and included 7 male to mixed-sex flocks and 3 female to mixed-sex flocks sampled 3 times at 6 day intervals.

The rate at which individuals move from exposed to infectious states (*σ*_*i*_), recovery rate (*γ*_*i*_), and mortality rate (*μ*_*i*_) of sex *i* were estimated from the experimental canary infections described above. Canaries are a common model for house finch-MG infections and have similar pathology and susceptibility to MG as house finches (Hawley et al., 2011). Parameter *σ*_*i*_ is calculated as the inverse of the average number of days from exposure until presentation of symptomatic disease (i.e. eye score < 0) while *γ*_*i*_ is the inverse of the average number of days from exposure until recovery from symptomatic disease (i.e. eye score = 0). Mortality rate (*μ*_*i*_) is calculated as the daily rate which individuals die after exposure and is given by 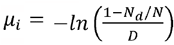 where *N*_*d*_ /*N* is the proportion of individuals that died during the duration of the experiment (*D*). We also calculated the basic reproductive number, given by *R*_o_ = *β*_*ii*_ / *β*_*i*_, for all male and all female populations (Anderson & May, 1986). See Table 1 for full list of parameter definitions and their values.

**Table 1.** Description of SEIR model parameters and their associated symbols and values.

|  |  |  |
| --- | --- | --- |
| Model parameter name | Symbol | Value |
| Transmission rate from male to male | $\beta_{MM}$ | 0.335 |
| Transmission rate from male to 50:50 mixed flock | $\beta_{ME}$ | 0.134 |
| Transmission rate from female to female | $\beta_{FF}$ | 0.138 |
| Transmission rate from female to 50:50 mixed flock | $\beta_{FE}$ | 0.128 |
| Rate at which males move from exposed to infectious states | $\sigma_M$ | 0.273 |
| Rate at which females move from exposed to infectious states | $\sigma_F$ | 0.200 |
| Recovery rate of male individuals | $\gamma_M$ | 0.048 |
| Recovery rate of female individuals | $\gamma_F$ | 0.091 |
| Daily mortality rate of male individuals | $\mu_M$ | 0.016 |
| Daily mortality rate of female individuals | $\mu_F$ | 0.005 |

To determine the robustness of our parameter calculations we conducted a sensitivity analysis for all parameters except for mortality rate. We ran simulations of the previously described SEIR model that individually varied each parameter by ± 1.0 and ± 0.5 SD (see Supplemental methods and the supplemental sensitivity analysis database for more information). In all simulations, epidemic size increased with the number of male birds in the flock. All-female flock epidemics were largest at *β*_*ff*_ +1 SD, though the epidemic was still lower than the smallest all-male flock epidemic which occurred at *γ*_*m*_ -1 SD (64% vs. 77% of individuals infected, respectively). Only one other all-female flock reached an epidemic size where >11% of individuals were infected (*β*_*ff*_ +0.5 SD; 30%) and no other all-male flocks had epidemics where <87% of the flock was infected.

## Results

### Experimental results

MG-exposed male canaries had higher total eye scores (*β(male)*=1.85 ± 0.22 SE, *t*=8.49, *p*<0.001; Figure 2A & Table S2) and pathogen loads (*β(male)* =1.34 ± 2.04 SE, χ^2^ =3.68, *df*=1, *p*=0.06; Figure 2B & Table S3) than MG-exposed females. In general, females and control birds maintained slightly higher body fat over time (sex*day: *β(male)* =0.02 ± 0.01 SE, χ^2^=7.48, *df*=1, *p*<0.01; MG-exposure*day: *β(mg)* =-0.01 ± 0.01 SE, χ^2^=3.81, *df*=1, *p*=0.05; Figure S1A & Table S4) and had greater mass (main effect of sex: *β(male)* =-5.09 ± 2.23 SE, χ^2^=3.81, *df*=1, *p*=0.05; main effect of MG-exposure: *β(mg)* =-4.57 ± 2.35 SE, χ^2^=4.04, *df*=1, *p*=0.04; Figure S1B & Table S5). However, there was no interactive effect of sex and treatment or sex, treatment, and time on body fat or mass. MG-exposed males produced more MG-specific antibodies over time than females (sex*day: *β(male)* =0.003 ± 0.001 SE, χ^2^=8.36, *df*=1, *p*<0.01; Figure 2C & Table S6). Further, there was a significant interaction between sex and MG-exposure on the change in relative abundance of monocytes after exposure resulting in MG-exposed males having a greater increase in monocyte relative abundance than all other groups (sex*treatment: *SS*=78.67, *F*_*1,17*_=6.91, *p*=0.02; Figure S2 & Table S7). Finally, male canaries generally had higher hematocrit levels and a greater increase in the relative abundance of eosinophils regardless of exposure status (sex: *β(male)* =4.87 ± 2.22 SE, χ^2^=14.71, *df*=1, *p*<0.001; *SS*=284.73, *F*_*1,17*_=6.13, *p*=0.02; Figure S3 & Tables S7 & S8). In contrast, relative abundance of eosinophils prior to exposure were higher in females than in males (*SS*=291.37, *F*_*1,20*_=3.21, *p*<0.01; Table S9).

**Figure 2.**
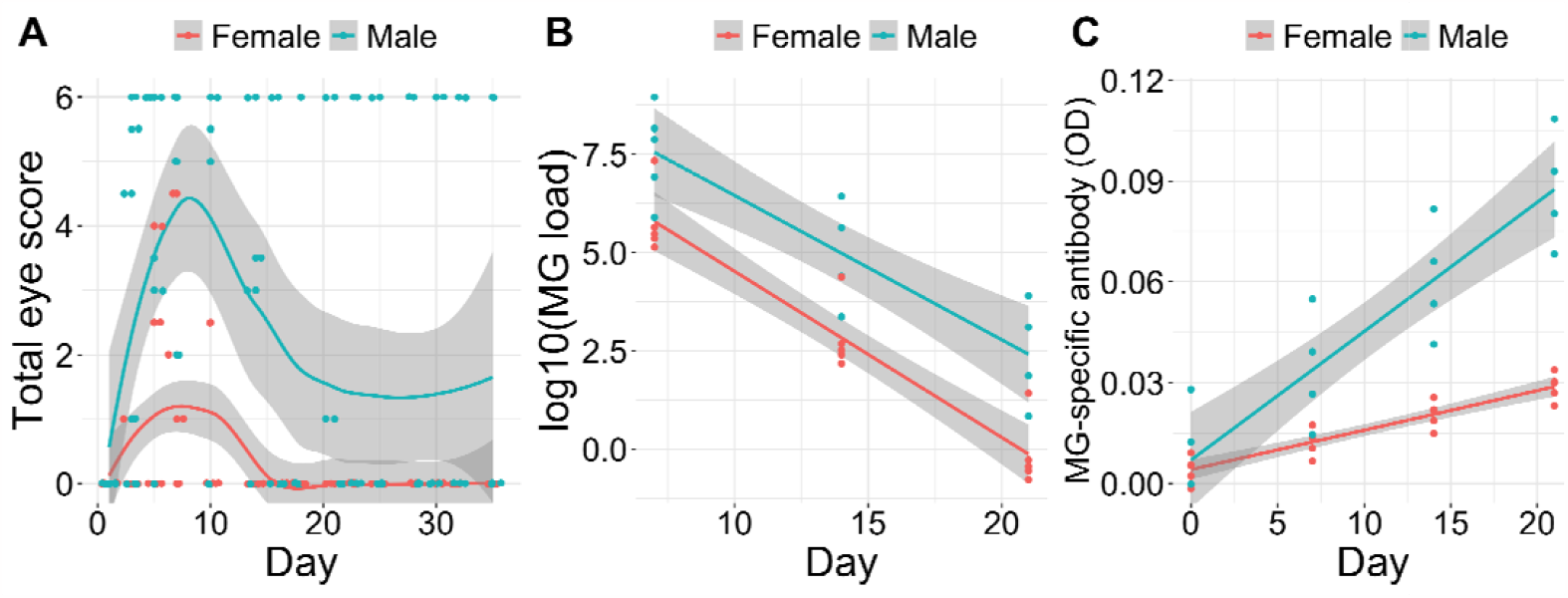
Differential effect of canary sex (*Serinus canaria domestica*) on **A**) total eye score (*β*=1.85, *t*=8.49, *p*<0.001), **B**) log-transformed pathogen load (χ^2^=3.68, *df*=1, *p*=0.06), and **C**) MG-specific antibody levels (χ^2^=6.44, *df*=1, *p*=0.01) after exposure to *Mycoplasma gallisepticum* (MG). Points in (**A**) are raw data. Points in (**B**) & (**C**) are predicted model values from linear mixed effects models that examined the main effects and interactions of sex, MG-exposure status, and time since infection on pathogen load and antibody level, respectively. Gray shading represents associated 95% confidence bands.

### SEIR model results

Epidemic size drastically increased with the proportion of the population that is male (Figure 3). The male-only population experienced a large and rapid epidemic, with 99% of individuals becoming exposed within 60 days (i.e. sum of individuals in the EIRM classes) and 100% of individuals becoming infected during the epidemic (i.e. sum of individuals in the IRM classes at the end of the simulation; *R*_o_=7.04; Figure 3A). In contrast, the female-only population experienced a small outbreak that fizzled out, with 10% of individuals becoming infected in 60 days (*R*_o_= 1.51; Figure 3E). However, shifting from an all-female population to a population where only 25% were male resulted in a 636% increase in epidemic size (shift from 10.29 to 75.41% total prevalence; Figure 3D & E). Further, the 50:50 population experienced a large epidemic, a total of 89% becoming infected during the entire epidemic (Figure 3C). Additionally, the simulated 50:50 epidemic was slightly male-biased, with 96% of males becoming infected during the epidemic compared to 82% of females.

**Figure 3.**
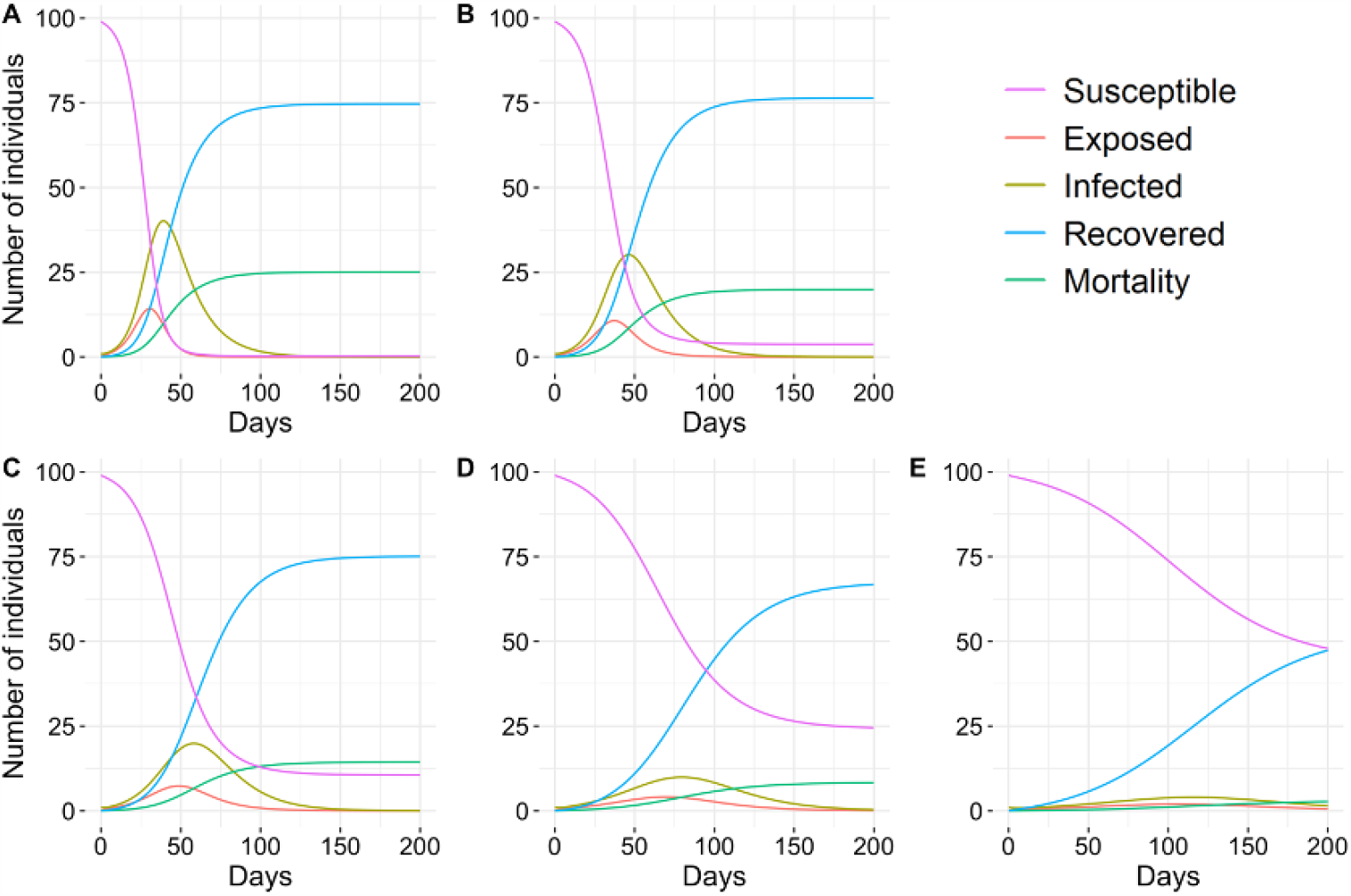
Results of the Susceptible, Exposed, Infected, Recovered, Mortality transmission model when the initial proportion of susceptible individuals that were male was **A**) 100%, **B**) 75%, **C**) 50%, **D**) 25%, and **E**) 0% (i.e. 100% female). Starting conditions were *S*=99, *E*=0, *I*=1, *R*=0, *M*=0. In all models with male individuals, the initial infected bird was male.

Finally, to determine the effect of physiology on transmission, we manipulated parameters measured in isolation, independent from behaviour (i.e., latency period and recovery rate) in the male only population. First, we found that decreasing *γ* (recovery rate) in males to the female rate resulted in *R*_o_ being nearly halved (*R*_o_= 3.69) but still high enough to cause an epidemic as *R*_o_> 1. Second, we found that decreasing male *γ* and *σ* (rate at which individuals move from exposed to infectious states) to female rates in a simulated epidemic of the 50:50 population more than halved peak prevalence (shift from 20 to 9%). These parameter changes lead to epidemic dynamics in the 50:50 population that closely resemble those of the 25% male population.

## Discussion

Here we set out to examine the previously unexplored potential for male-biased pathology, independent of behavior, to affect transmission using an avian host-pathogen system. We built a sex-dependent transmission model parameterized with isolated, individual-based experimental exposures and experimental transmission data. Our controlled MG-exposure experiment demonstrated that male canaries have shorter incubation periods, longer recovery periods, and higher pathogen burdens than females. Using a multistate transmission model parameterized with exposure-controlled pathological data of domestic canaries and transmission rates from house finch flocks of varying sex-ratios, we show that epidemic size rapidly increases with the proportion of male birds in a flock. These results lend support to the hypothesis that there are epidemiological consequences to sex-biased resistance and pathology. This study is the first to model sex-biased transmission in this system, and is one of few to model sex-biased transmission dynamics in an avian pathogen system (Grear et al., 2009; Lachish et al., 2011; Mougeot et al., 2005; Zuk & McKean, 1996).

Isolated experimental exposures in canaries suggest that male bird physiology is contributing to MG transmission dynamics independent of sex-biased behavioural differences. Male incubation periods are 27% faster and they remain infectious for 48% longer than females, meaning the period which they can transmit disease is nearly twice as long. Similarly, male house finch incubation periods were 30% faster in Adelman et al., 2015. Further, male birds had higher pathogen loads and greater pathology than females, suggesting male canaries would deposit greater quantities of pathogen on fomites, a critical transmission mode in this system (Adelman et al., 2015; Dhondt et al., 2007). We also found that male-biased mortality resulted in a 0.79 males per female ratio in our 50:50 flock simulation. Surveys of house finch populations shortly after the initial introduction of MG found similar effects with sex ratios shifting from even or slightly male-biased to female biased (Nolan et al., 1998). Given the continued abundance of MG in house finch populations, it is possible that sex-biased mortality may have caused long lasting and widespread shifts in adult sex ratio or effects on maternal manipulation of embryo sex ratios (Badyaev et al., 2006; DuRant et al., 2016). However, surveys of pre- and post-epidemic house finch sex-ratios are needed to further understand the effects of sex-biased mortality on populations.

While there is limited available data on sex-biased pathology in the house finch-MG system, there is some supporting evidence that male house finches have significantly greater pathology (i.e. eye inflammation) than females (Adelman et al., 2015; Hawley et al., 2007; Moyers et al., 2018). However, the evidence is inconsistent as some studies have found greater pathology in females (Altizer et al., 2004; Leon & Hawley, 2017) and many find no difference between sexes (Sydenstricker et al., 2006; Thomason et al., 2017; Vinkler et al., 2018; Weitzman et al., 2021). Robust studies explicitly testing for sex-biased resistance in house finch populations are lacking and needed to better understand these dynamics. Experiments on juvenile birds could be especially informative, as this age class is the largest group of naïve individuals and differential pathology or behavior in the class remains unexplored. The transmission model simulations that varied population sex-ratio supported male-biased MG transmission. Epidemic size drastically increased with the proportion of birds that were male, increasing by 872% from the all-female to the all-male simulation. Further, peak prevalence more than halved when we simulated 50:50 epidemics where male recovery and incubation rates were equivalent to female rates, providing further evidence that lower resistance in males is largely driving sex-biased transmission dynamics. Our single-season model assumes full immunity after infection, but it is possible that sex-biased differences in loss of immunity may exist as well, potentially exacerbating the sex-bias we find in the study.

The mechanism driving female-biased resistance (ability to prevent or limit pathogen growth) in canaries is unclear though we did find that prior to exposure, females had higher levels of circulating eosinophils than males. While the function of eosinophils in birds is unclear, they have been associated with greater body condition and parasite resistance in passerines (Deem et al., 2011; Owen et al., 2013; Owen & Moore, 2008). After exposure, MG exposed males produced more MG-specific antibodies and increased eosinophils to levels equal to those females had prior to exposure. This larger induced immune response in males is likely a response to their higher pathogen burdens as the larger response did not translate to faster recovery times or lower MG loads. This result supports prior literature on female immunocompetence (Zuk & McKean, 1996) as females were able to resist MG growth, resulting in lower pathology and faster recovery. Ultimately, further research is needed to reveal the proximate mechanisms that support greater immunocompetence in females than males.

Using avian-MG host pathogen systems we found that male-biased transmission can, in part, be attributed to male-biased pathology. Sex, and thus sex-ratio of the susceptible population, can be an important source of heterogeneity in epidemics. However, direct assessment of sex-biased transmission driven by sex-based differences in pathology in wildlife is largely unexplored with most studies focusing on behavioural drivers of transmission. Here, we demonstrate that sex-biased pathology, independent of behaviour, can have large impacts on transmission dynamics. While the role of sex-biased immunity is often difficult to disentangle from behaviour, epidemic models parameterized by experimentally controlled exposures provide an opportunity to better understand how aspects of pathology contribute to transmission.

## Supporting information

Sensitivity analysis

## Acknowledgments

We thank James Adelman for providing helpful comments and discussion and two anonymous reviewers. Funds were provided by grants to S.E.D. from the National Science Foundation (1941861) and Arkansas Biosciences Institute. Canary experimental methods were approved by the University of Arkansas International Animal Care and Use Committee.

