## Supplemental information for "Male pathology regardless of behaviour drives transmission in an avian host-pathogen system"

Supplemental Methods and Results for: **Male pathology regardless of behaviour drives transmission in an avian host-pathogen system**

**Authors:** Erin L. Sauer^1^, Chloe Connelly^1^, Weston Perrine^1^, Ashely C. Love^1,2^, Sarah E. DuRant^1^

**Affiliations:**

^1^Department of Biological Sciences, University of Arkansas, Fayetteville, AR.

^2^Ecology and Evolutionary Biology, University of Connecticut, Storrs, CT.

**Supplemental Methods**

*House finch data analysis –* Using the published house finch raw data from Adelman et al. 2015, we tested for the effect of sex on total eye score, we conducted a Poisson distributed generalized additive mixed model that included a smoothing spline to model non-linear effects over time identical to the one used for the canary data. Similar to the canary results, MG-exposed male house finches had higher total eye scores (*β(male)*=0.79 ± 0.27 SE, *t*=2.92, *p*=0.004, s(Day): *edf*=1.98, *F*=19.85, *p*<0.001) than MG-exposed females.

*Canary exposures* – The canary exposure data used to parameterize the SEIR (susceptible, exposed, infected, removed) model came from an unrelated forthcoming study designed to test changes in feeding behaviors during MG infection.

*SEIR model with frequency-dependent transmission* – Frequency-dependent transmission dynamics are given by the system of differential equations:

$$\frac{dS_{i}}{dt}=-\sum_{j} \beta_{ij}I_{j}S_{i}$$

$$\frac{dE_{i}}{dt}=\sum_{j} \beta_{ij}I_{j}S_{i}-\sigma_{i}E_{i}$$

$$\frac{dI_{i}}{dt}=\sigma_{i}E_{i}-\gamma_{i}I_{i}-\mu_{i}I_{i}$$

$$\frac{dR_{i}}{dt}=\gamma_{i}I_{i}$$

$$\frac{dM_{i}}{dt}=\mu_{i}I_{i}$$

*Transmission coefficients* – Transmission rate from infectious sex *i* to susceptible host type *j* (*β_ij_*) was estimated from two published transmission datasets (Adelman *et al.* 2015; Moyers *et al.* 2018) and is given by: $\beta_{ij}=\tau\bar{c}$ . Where $\tau$ is the prevalence of new infections among susceptible individuals at that sampling period and $\bar{c}$ is the contact rate or the number of individuals not previously infected divided by the duration of time since the last sampling period (Anderson & May 1986). To calculate flock-level transmission coefficients, we averaged across multiple sampling periods. Final *β_ij_* values are the average of multiple same-type flock-level transmission rates. An individual was considered infected when MG load was > 100 gene copies. Table S1 shows sample point-level transmission coefficients. See Adelman *et al.* (2015) and Moyers *et al.* (2018) for detailed information on experimental design, data collection, and results.

*Sensitivity analysis* – To determine the robustness of our parameter calculations we conducted a sensitivity analysis for all parameters except for mortality rate. We ran simulations of the SEIR model described in the main text that individually varied each parameter by ± 1.0 and ± 0.5 SD. This resulted in five simulations of varying sex-ratios for 33 different parameter sets. In all simulations, epidemic size increased with the number of male birds in the flock. The detailed results of this analysis are available in the sensitivity analysis database.

| Table S1 \| House finch (*Haemorhous mexicanus*) flock-level *Mycoplasma gallisepticum* transmission coefficients calculated from publicly available datasets (Adelman *et al.* 2015; Moyers *et al.* 2018) for every sampling time point. | | | | | | |
| --- | --- | --- | --- | --- | --- | --- |
| **Female only flocks** | | | | | | |
|  | Sample day | | | | |  |
| Flock ID | 4 | 8 | 12 | 16 | 20 | Mean flock β |
| 1 | 0.000 | 0.667 | 0.000 | 1.000 | - | 0.417 |
| 2 | 0.000 | 0.000 | 0.333 | 0.000 | 0.000 | 0.067 |
| 4 | 0.000 | 0.000 | 0.333 | 0.000 | 0.000 | 0.067 |
| 5 | 0.000 | 0.000 | 0.000 | 0.000 | 0.000 | 0.000 |
| **Male only flocks** | | | | | | |
|  | Sample day | | | | |  |
| Flock ID | 4 | 8 | 12 | 16 | 16 | Mean flock β |
| 6 | 0.000 | 0.000 | 0.667 | 0.000 | 0.000 | 0.133 |
| 7 | 0.000 | 0.667 | 1.000 |  |  | 0.556 |
| 8 | 0.000 | 0.333 | 0.000 | 0.000 | 0.000 | 0.067 |
| 9 | 0.000 | 0.333 | 1.000 |  |  | 0.444 |
| 10 | 0.000 | 0.333 | 0.500 | 0.000 | 1.000 | 0.367 |
| 11 | 0.000 | 0.333 | 1.000 |  |  | 0.444 |
| **Female index to mixed-sex flock** | | | | | | |
|  | Sample day | | |  | | |
| Flock ID | 6 | 12 | 18 | Mean flock β | | |
| 1 | 0.000 | 0.167 | 0.000 | 0.056 | | |
| 9 | 0.167 | 0.381 | 0.267 | 0.267 | | |
| 10 | 0.167 | 0.000 | 0.000 | 0.056 | | |
| **Male index to mixed-sex flock** | | | | | | |
|  | Sample day | | |  | | |
| Flock ID | 6 | 12 | 18 | Mean flock β | | |
| 2 | 0.167 | 0.000 | 0.375 | 0.181 | | |
| 3 | 0.167 | 0.000 | 0.000 | 0.056 | | |
| 4 | 0.000 | 0.500 | 0.000 | 0.167 | | |
| 5 | 0.167 | 0.563 | 0.000 | 0.243 | | |
| 6 | 0.167 | 0.000 | 0.000 | 0.056 | | |
| 7 | 0.167 | 0.375 | 0.000 | 0.181 | | |
| 8 | 0.000 | 0.167 | 0.000 | 0.056 | | |

**Supplemental Results**

Figure S1 | Differential effect of canary sex (*Serinus canaria domestica*) and *Mycoplasma gallisepticum* (MG) exposure status on **A**) fat score and **B**) body mass (g). Points are predicted model values from linear mixed effects models that examined the main effects and interactions of sex, MG-exposure status, and days since infection on **A**) fat score and **B**) body mass. Gray shading represents associated 95% confidence bands.

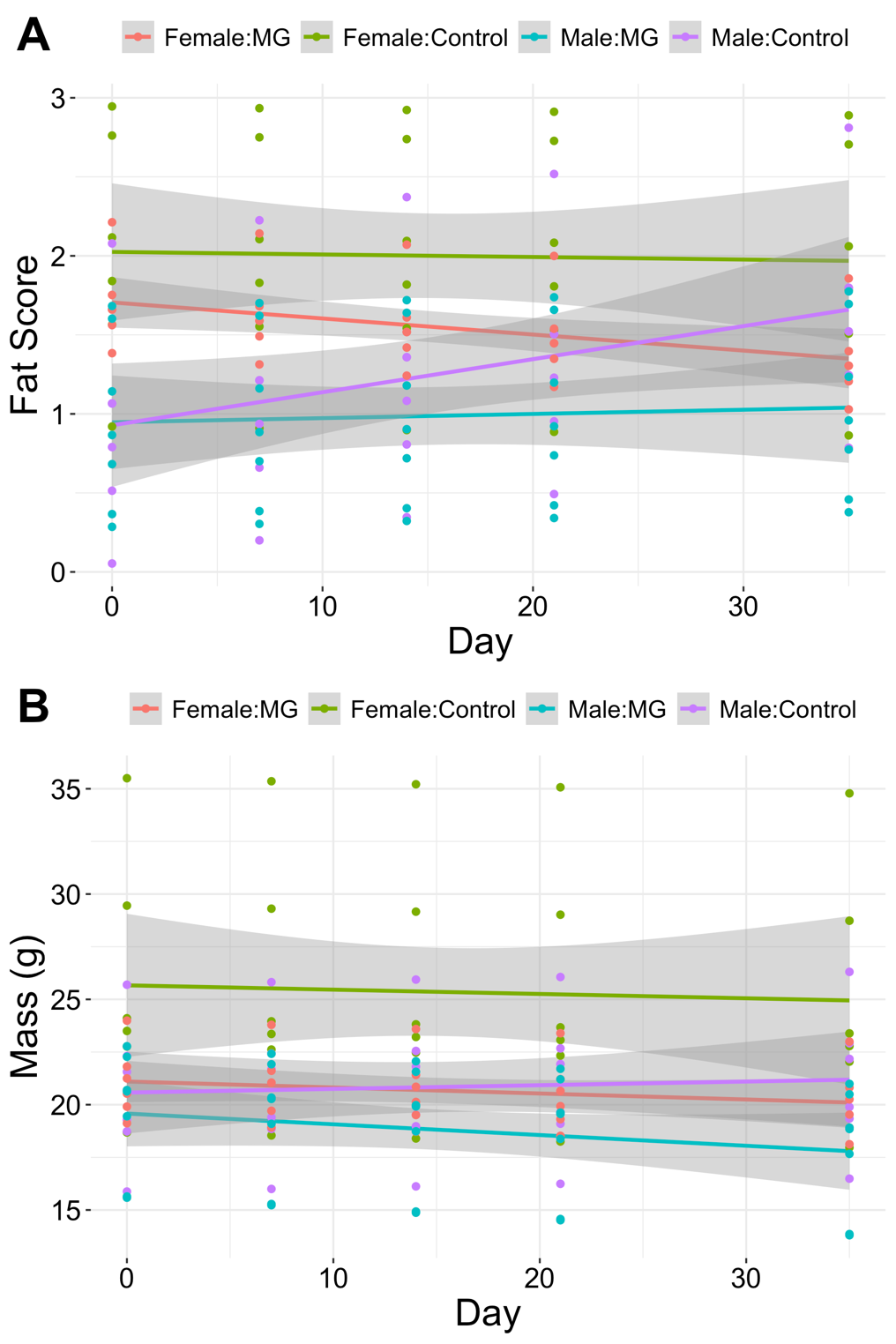

Figure S2 | Box plot showing the change in relative abundance of white blood cells in canaries (*Serinus canaria domestica*) one week after *Mycoplasma gallisepticum* (MG) or sham (control) exposure in male and female birds.

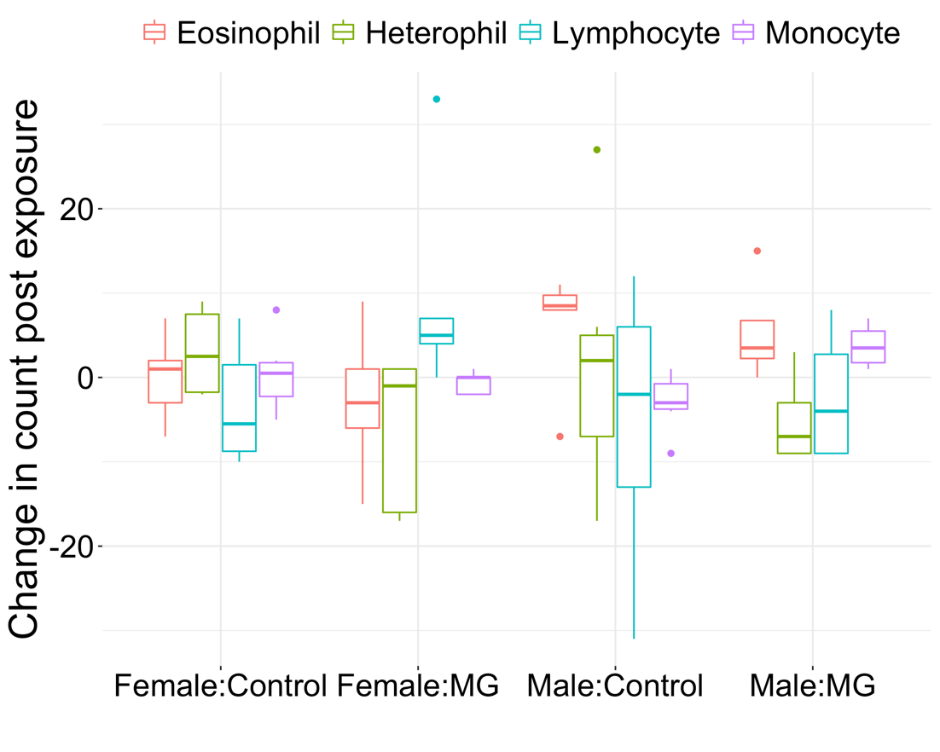

Figure S3 | Differential effect of canary sex (*Serinus canaria domestica*) and *Mycoplasma gallisepticum* (MG) exposure status on hematocrit %. Predicted model values from a linear mixed effects model that examined the main effects and interactions of sex, MG-exposure status, and days since infection on hematocrit %. Gray shading represents associated 95% confidence bands.

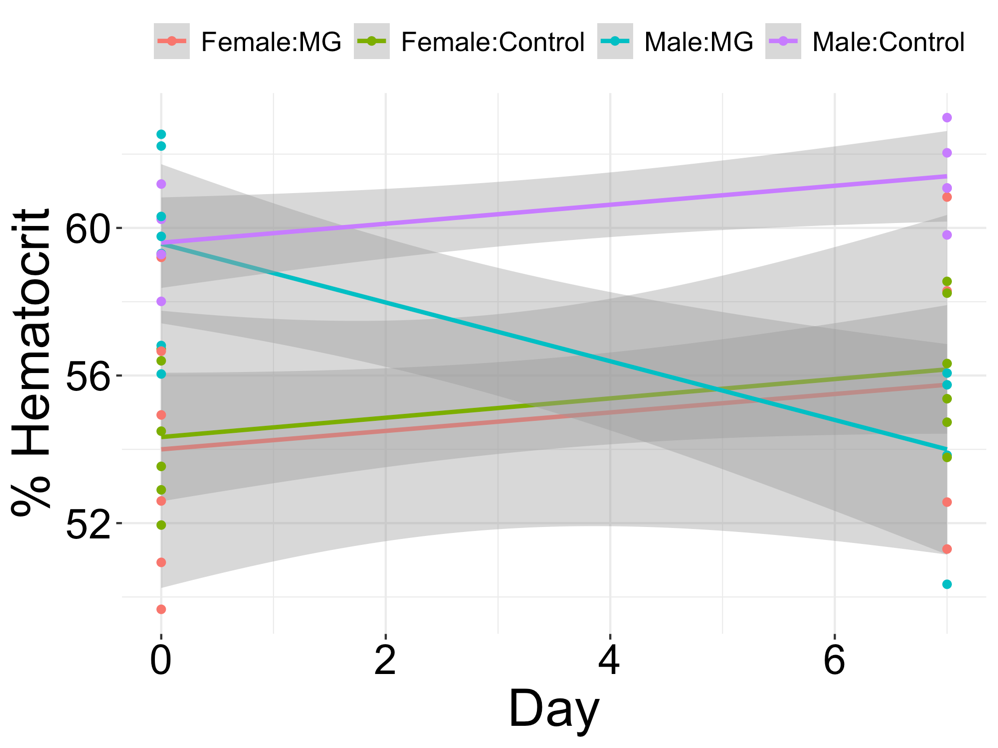

Figure S4 | Results of the frequency-dependent Susceptible, Exposed, Infected, Recovered, Mortality transmission model when the initial proportion of susceptible individuals that were male was **A**) 100%, **B**) 75%, **C**) 50%, **D**) 25%, and **E**) 0% (i.e. 100% female). Starting conditions were *S*=99, *E*=0, *I*=1, *R*=0, *M*=0. In all models with male individuals, the initial infected bird was male.

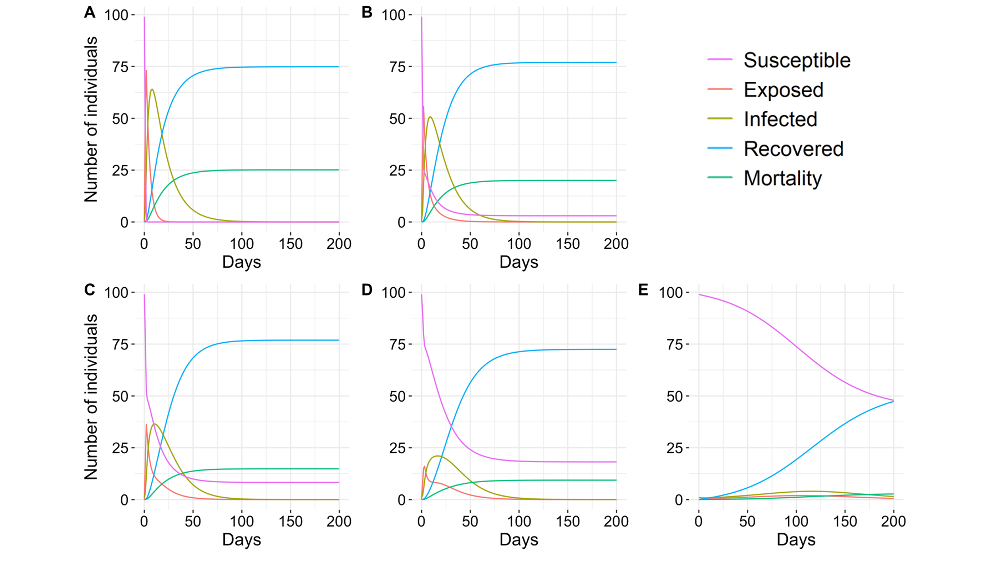

Tables

| Table S2 \| Results from a Poisson-distributed generalized additive mixed model examining the effect of canary sex (*Serinus canaria domestica*) and the smoothed term time since *Mycoplasma gallisepticum* (MG) exposure on conjunctival inflammation (total eye score). | | | | |
| --- | --- | --- | --- | --- |
| Parametric coefficients |  |  |  |  |
|  | Coefficient | SE | t value | p value |
| Sex:Male | 1.8499 | 0.218 | 8.486 | <0.001 |
| Approximate significance of smooth terms: |  |  |  |  |
|  |  | edf | F statistic | p value |
| s(Day) |  | 2.883 | 10.72 | <0.001 |

| Table S3 \| Results from the linear mixed-effects model examining the effect of canary sex (Serinus canaria domestica), days since Mycoplasma gallisepticum (MG) exposure, and their interaction on log10-transformed MG load and ANOVA. | | | |
| --- | --- | --- | --- |
| Linear mixed effects model results | | | |
|  | Coefficient | SE | t value |
| Intercept | 8.744 | 1.425 | 6.135 |
| SexMale | 1.343 | 2.039 | 0.659 |
| Day | -0.422 | 0.085 | -4.963 |
| SexMale:Day | 0.061 | 0.125 | 0.490 |
| ANOVA results | | | |
|  | Chi squared | df | p value |
| Sex | 3.677 | 1 | 0.055 |
| Day | 39.816 | 1 | <0.001 |
| Sex:Day | 0.240 | 1 | 0.624 |

| Table S4 \| Results from the linear mixed-effects model examining the effects of canary sex (Serinus canaria domestica), days since Mycoplasma gallisepticum (MG) or sham exposure, MG exposure status, and their interactions on fat reserve score and ANOVA. | | | |
| --- | --- | --- | --- |
| Linear mixed effects model results | | | |
|  | Coefficient | SE | t value |
| Intercept | 2.025 | 0.286 | 7.087 |
| SexMale | -1.097 | 0.404 | -2.715 |
| Day | -0.002 | 0.006 | -0.256 |
| TreatmentMG | -0.319 | 0.407 | -0.786 |
| SexMale:Day | 0.023 | 0.009 | 2.532 |
| SexMale:TreatmentMG | 0.338 | 0.566 | 0.598 |
| Day:TreatmentMG | -0.009 | 0.009 | -0.921 |
| SexMale:Day:TreatmentMG | -0.010 | 0.014 | -0.720 |
| ANOVA results | | | |
|  | Chi squared | df | p value |
| Sex | 6.426 | 1 | 0.011 |
| Day | 1.121 | 1 | 0.290 |
| Treatment | 1.429 | 1 | 0.232 |
| Sex:Day | 7.483 | 1 | 0.006 |
| Sex:Treatment | 0.152 | 1 | 0.696 |
| Day:Treatment | 3.805 | 1 | 0.051 |
| Sex:Day:Treatment | 0.518 | 1 | 0.472 |

| Table S5 \| Results from the linear mixed-effects model examining the effects of canary sex (*Serinus canaria domestica*), days since *Mycoplasma gallisepticum* (MG) or sham exposure, MG exposure status, and their interactions on host body mass and ANOVA. | | | |
| --- | --- | --- | --- |
| Linear mixed effects model results | | | |
|  | Coefficient | SE | t value |
| (Intercept) | 25.667 | 1.657 | 15.492 |
| SexMale | -5.091 | 2.343 | -2.173 |
| Day | -0.020 | 0.025 | -0.814 |
| TreatmentMG | -4.566 | 2.350 | -1.943 |
| SexMale:Day | 0.038 | 0.036 | 1.066 |
| SexMale:TreatmentMG | 3.570 | 3.268 | 1.092 |
| Day:TreatmentMG | -0.008 | 0.037 | -0.214 |
| SexMale:Day:TreatmentMG | -0.060 | 0.054 | -1.115 |
| ANOVA results | | | |
|  | Chi squared | df | p value |
| Sex | 3.812 | 1 | 0.051 |
| Day | 1.715 | 1 | 0.190 |
| Treatment | 4.039 | 1 | 0.044 |
| Sex:Day | 0.197 | 1 | 0.657 |
| Sex:Treatment | 0.760 | 1 | 0.383 |
| Day:Treatment | 1.812 | 1 | 0.178 |
| Sex:Day:Treatment | 1.243 | 1 | 0.265 |

| Table S6 \| Results from the linear mixed-effects model examining the effect of canary sex (Serinus canaria domestica), days since Mycoplasma gallisepticum (MG) exposure, and their interaction on MG specific antibody levels (optical density) and ANOVA. | | | |
| --- | --- | --- | --- |
| Linear mixed effects model results | | | |
|  | Coefficient | SE | t value |
| Intercept | 0.004 | 0.010 | 0.403 |
| SexMale | 0.003 | 0.015 | 0.182 |
| Day | 0.001 | 0.001 | 1.916 |
| SexMale:Day | 0.003 | 0.001 | 2.891 |
| ANOVA results | | | |
|  | Chi squared | df | p value |
| Sex | 6.436 | 1 | 0.011 |
| Day | 26.584 | 1 | <0.001 |
| Sex:Day | 8.357 | 1 | 0.004 |

| Table S7 \| MANOVA examining the effect of canary sex (*Serinus canaria domestica*) and *Mycoplasma gallisepticum* (MG) or sham exposure on the change in relative abundance of leukocytes from pre-exposure to one week post exposure. | | | | |
| --- | --- | --- | --- | --- |
| MANOVA | | | | |
|  | Pillai's trace | df | F value | p value |
| Sex | 0.294 | 1,14 | 1.455 | 0.268 |
| Treatment | 0.308 | 1,14 | 1.558 | 0.240 |
| Sex:Treatment | 0.351 | 1,14 | 1.889 | 0.168 |
| Residuals |  | 17 |  |  |
| Monocyte | | | | |
|  | Sum Sq | df | F value | p value |
| Sex | 0.471 | 1 | 0.041 | 0.841 |
| Treatment | 33.983 | 1 | 2.986 | 0.102 |
| Sex:Treatment | 78.667 | 1 | 6.913 | 0.018 |
| Residuals | 193.450 | 17 |  |  |
| Eosinophil | | | | |
| Sex | 284.730 | 1 | 6.133 | 0.024 |
| Treatment | 19.650 | 1 | 0.423 | 0.524 |
| Sex:Treatment | 4.140 | 1 | 0.089 | 0.769 |
| Residuals | 789.300 | 17 |  |  |
| Heterophil | | | | |
| Sex | 0.390 | 1 | 0.004 | 0.951 |
| Treatment | 338.110 | 1 | 3.341 | 0.085 |
| Sex:Treatment | 9.540 | 1 | 0.094 | 0.763 |
| Residuals | 1720.530 | 17 |  |  |
| Lymphocyte | | | | |
| Sex | 230.690 | 1 | 1.613 | 0.221 |
| Treatment | 357.580 | 1 | 2.500 | 0.132 |
| Sex:Treatment | 133.250 | 1 | 0.932 | 0.348 |
| Residuals | 2431.720 | 17 |  |  |

| Table S8 \| Results of multiple linear mixed-effect models examining the effects of canary sex (Serinus canaria domestica), days since Mycoplasma gallisepticum (MG) or sham exposure, MG exposure status, and their interactions on % hematocrit and ANOVA. | | | |
| --- | --- | --- | --- |
| Linear mixed effects model results | | | |
|  | Coefficient | SE | t value |
| Intercept | 54.850 | 1.498 | 36.614 |
| SexMale | 4.870 | 2.222 | 2.192 |
| Day | 0.014 | 0.099 | 0.144 |
| TreatmentMG | -0.864 | 2.166 | -0.399 |
| SexMale:Day | 0.074 | 0.148 | 0.503 |
| SexMale:TreatmentMG | -0.889 | 3.088 | -0.288 |
| Day:TreatmentMG | -0.004 | 0.150 | -0.026 |
| SexMale:Day:TreatmentMG | -0.182 | 0.216 | -0.842 |
| ANOVA results | | | |
|  | Chi squared | df | p value |
| Sex | 14.705 | 1 | <0.001 |
| Day | 0.011 | 1 | 0.917 |
| Treatment | 3.355 | 1 | 0.067 |
| Sex:Day | 0.009 | 1 | 0.923 |
| Sex:Treatment | 1.283 | 1 | 0.257 |
| Day:Treatment | 0.711 | 1 | 0.399 |
| Sex:Day:Treatment | 0.709 | 1 | 0.400 |

| Table S9 \| MANOVA examining the effect of canary sex (*Serinus canaria domestica*) on the relative abundance of leukocytes prior to *Mycoplasma gallisepticum* (MG) or sham exposure. | | | | |
| --- | --- | --- | --- | --- |
| MANOVA | | | | |
|  | Pillai's trace | df | F value | p value |
| Sex | 0.391 | 1,20 | 3.205 | 0.035 |
| Residuals |  | 23 |  |  |
| Monocyte | | | | |
|  | Sum Sq | df | F value | p value |
| Sex | 12.523 | 1 | 1.247 | 0.276 |
| Residuals | 230.917 | 23 |  |  |
| Eosinophil | | | | |
| Sex | 291.370 | 1 | 8.176 | 0.009 |
| Residuals | 819.670 | 23 |  |  |
| Heterophil | | | | |
| Sex | 26.420 | 1 | 0.285 | 0.599 |
| Residuals | 2131.020 | 23 |  |  |
| Lymphocyte | | | | |
| Sex | 112.700 | 1 | 0.355 | 0.557 |
| Residuals | 7300.200 | 23 |  |  |
